# Measurement and classification of bold-shy behaviours in medaka fish

**DOI:** 10.1101/2024.10.18.618696

**Authors:** Saul Pierotti, Ian Brettell, Tomas Fitzgerald, Cathrin Herder, Narendar Aadepu, Christian Pylatiuk, Joachim Wittbrodt, Ewan Birney, Felix Loosli

## Abstract

**Motivation:** Boldness-shyness is considered a fundamental axis of behavioural variation in humans and other species, with obvious adaptive causes and evolutionary implications. Besides an individual’s own genetics, this phenotype is also affected by the genetic makeup of peers in the individual’s social environment. To identify genetic determinants of variation along the bold-shy behavioural axis, a reliable experimental and analytical setup able to highlight direct and indirect genetic effects is needed.

**Results:** We describe a custom assay designed to detect bold-shy behaviours in medaka fish, combining an open-field and novel-object component. We use this assay to explore direct and social genetic effects on the behaviours of 307 pairs of fish from five inbred medaka strains. Applying a Hidden Markov Model (HMM) to classify behavioural modes, we find that direct genetic effects influence the proportions of time the five strains spent in slow-moving states, explaining up to 29.7% of the variance in time spent in those states. We also found that an individual’s behaviour is influenced by the genetics of its tank partner, explaining up to 8.64% of the variance in the time spent in slow-moving states. Our behavioural assay in combination with the HMM analysis is applicable to follow-up genetic linkage studies of genetic variants involved in direct behavioural effects and indirect social genetic effects. A suitable genetic resource for such studies, the Medaka Inbred Kiyosu-Karlsruhe panel (MIKK) has recently been established.

## Introduction

Animals modulate their behaviour to obtain food and other resources, particularly mates, while minimising the risk of predation or other threats. Many animals exhibit complex behaviours which balance the different rewards and penalties that might be present in their environment (Altendorf *et al*., 2001; Higginson *et al*., 2012). Furthermore, animals which congregate in social groups can react to the behaviour of other individuals of the same species and adapt to the social environment (Ward and Webster, 2016). These social interactions may improve the overall ability of the group to detect predation (Kelley *et al*., 2003) and place animals in proximity to conspecific individuals for better access to reproduction (Beck *et al*., 2021; Uzsák and Schal, 2013). As such, behaviour in general and social behaviour in particular are critical traits for the survival and reproduction of many species (Robinson *et al*., 2019).

Boldness-shyness is thought to be a fundamental axis of behavioural variation, with an obvious causal relationship to an individual’s likelihood of survival, and consequently with natural selection (Wilson *et al*., 1994). It represents an evolutionary trade-off between acquiring benefits (in terms of food or mates) and avoiding harms (in terms of predators or conspecific competitors), with each situation accompanied by its own optimal degree of risk (Lima and Dill, 1990). It is both heritable in many species (Svartberg, 2002; Culum Brown *et al*., 2007), and subject to change following different life experiences or under different environmental conditions (Culum Brown *et al*., 2007).

Two generic paradigms for measuring boldness include the “open field” test and the “novel object” test. The open field test involves observing the test subject while it moves freely in an unfamiliar experimental setting, and has been performed on many species including the bullfrog tadpole (Carlson and Langkilde, 2013), gecko (Nordberg *et al*., 2021), mouse (Garfield *et al*., 2011), African striped mouse (Yuen *et al*., 2017), rat (Baud *et al*., 2014), rabbit (Meijsser *et al*., 1989), common vole (Herde and Eccard, 2013), and Siberian dwarf hamster (Kanda *et al*., 2012). It is particularly favoured with fish, where the general interpretation is that shy individuals tend to react to novelty by reducing their activity and becoming more vigilant, whereas bold individuals show higher levels of activity and exploratory behaviour (C. Brown *et al*., 2007; Matsunaga and Watanabe, 2010; Dahlbom *et al*., 2011; Lucon-Xiccato and Bisazza, 2017; Lucon-Xiccato *et al*., 2020; Lucon-Xiccato, Loosli, *et al*., 2022; Alfonso *et al*., 2019; Hamilton *et al*., 2021). The second paradigm is the novel object test, where a novel object is introduced to the test subject’s environment. This test has also been used on many species including birds (Azevedo and Young, 2006), squirrels (Uchida *et al*., 2019), rabbits (Andersson *et al*., 2014), baboons (Carter *et al*., 2012), vervet monkeys (Blaszczyk, 2017), and grey mouse lemurs (Dammhahn and Almeling, 2012). Like the open field test, the novel object test has also been extensively used with fish (C. Brown *et al*., 2007; Hamilton *et al*., 2021; Schjolden *et al*., 2005; Wilson *et al*., 1993; Wright *et al*., 2006, 2003). Both the open field and novel object test also permit the measurement of habituation, where the response of the fish may change over time after growing accustomed to the new environment or object (Wong *et al*., 2010).

Environmental factors can have a profound influence on animal behaviour, and the social environment in particular can play a significant role in shaping behavioural phenotypes (Robinson, 1999). As well as numerous studies on mammals (Sachser *et al*., 2013; Snyder-Mackler *et al*., 2020), including humans (Kirschbaum *et al*., 1995), with complex social networks, lower vertebrates such as birds (Noguera *et al*., 2017), reptiles (Riley *et al*., 2017) and fish (Lucon-Xiccato, Montalbano, *et al*., 2022) have also been used for investigating the effect of an individual’s social environment on its behaviour.

As well as the environment, genetics also often impacts the traits of animals, including behaviour (Willoughby *et al*., 2023). The impact of genetic variation on behavioural traits, including the bold-shy paradigm, is well documented in a number of species (Oswald *et al*., 2013; Blanco *et al*., 2022; Kabelik *et al*., 2021; Bubac *et al*., 2021). As well as the direct genetic effects of variation on an individual’s behaviour, the indirect effect of genetic variation on the social peers of an individual has also been explored (Ribeiro *et al*., 2020; Chakrabarty *et al*., 2019; Baud *et al*., 2021; Anderson *et al*., 2017). This has been described as Indirect Genetic Effect (IGE) or Social Genetic Effect (SGE) (Moore *et al*., 1997; Baud *et al*., 2017).

The medaka fish (*Oryzias latipes*) is a long established model organism which has excellent genetics (relatively small genome, inbred lines), low husbandry costs and extensive genomic resources (Wittbrodt *et al*., 2002; Fitzgerald *et al*., 2022). It is a social animal and an established model to study social behaviour (Fukamachi *et al*., 2009; Imada *et al*., 2010; Kagawa, 2013; Ochiai *et al*., 2013; Nakayasu and Watanabe, 2013; Okuyama *et al*., 2014; Tsuboko *et al*., 2014; Yokoi *et al*., 2020, 2016, 2015; Isoe *et al*., 2016; Utagawa *et al*., 2016). To take advantage of the rich genetic and genomic resources offered by this model we established a joint open field and novel object test to examine the behaviour of different isogenic inbred medaka strains, with the aim of characterising variation along the boldness-shyness axis. To be able to study both direct and social genetic effects, we performed our assay on pairs of fish. Our aim was to determine: a) whether there were significant differences in bold-shy behaviours exhibited by five established inbred strains of medaka fish (*iCab, HdrR*, and *Ho5* from southern Japan, and *Kaga* and *HNI* from northern Japan); and b) whether there were significant differences in bold-shy behaviours exhibited by a given strain (*iCab*) dependent on the strain that it was partnered with. The former was intended to measure the effect of an individual’s own genes on its behaviour (a *direct* genetic effect), and the latter to measure the effect of the genes of the focal fish’s tank partner on the behaviour of the focal fish (*social* genetic effect). To ensure the robustness of the result we aimed for at least 47 biological replicates for each strain.

Our primary phenotyping scheme for the assay is a video recording of fish movement in a shallow tank of water (the shallow tank minimises the amount of depth variation). We chose to analyse this data using the well-established animal tracking software *idtracker*.*ai* (Romero-Ferrero *et al*., 2019) followed by a Hidden Markov Model (HMM) to infer behavioural modes from movement features obtained from the videos. HMMs are a simple and interpretable method for classifying stochastic, sequential observations as being generated by a set of hidden states (Mor *et al*., 2021). In this case, the HMM is used to extract the hidden behavioural states that produced the stochastic movement features that we observed. Our experimental design in combination with a Hidden Markov Model analysis of the recorded swim tracks allowed us to assess both direct and social effects on behaviour simultaneously, and to infer the degree to which variation in bold-shy behaviours is attributable to the differences in an individual’s own genetics, the differences in the genetics of their social companions, and stochastic variation.

With an experimental design of a small number of strains and a high number of replicates within a strain our approach can robustly measure overall genetic (whether direct or social) effects but is unable to pinpoint the responsible genetic loci. Using this approach, we can show that there are significant differences in hidden state occupancy across inbred medaka lines, likely linked to bold-shy behaviour. This work provides confidence in using the phenotyping and analysis scheme that we describe for studies focused on genomic locus discovery, which would require a different experimental design. The Medaka Inbred Kiyosu-Karlsruhe (MIKK) panel (Fitzgerald *et al*., 2022) will be a key resource for this future endeavour.

## Methods

### Fish husbandry

The inbred, isogenic medaka strains *iCab, HdrR*, and *Ho5* derived from the Southern Japanese medaka population and *HNI* and *Kaga* strains derived from the Northern Japanese medaka population were maintained as previously described (Loosli *et al*., 2000) – in closed stocks at the Institute of Biological and Chemical Systems, Biological Information Processing (IBCS-BIP), Karlsruhe Institute of Technology (KIT) in recirculatory systems, under 14 h light/10 h dark conditions at 26 °C . Fish husbandry was performed in accordance with EU directive 2010/63/EU guidelines as well as with German animal protection regulations (Tierschutzgesetz §11, Abs. 1, no. 1; Regierungspräsidium Karlsruhe, Germany; husbandry permits AZ35-9185.64/BH KIT). The fish facilities are under the supervision of the Regierungspräsidium Karlsruhe, who approved the experimental procedures. All strains were reared in 6-litre stocking tank systems. All fish used for the experiments were 6 months old and randomly selected from the stocking tanks prior to the experimental observation. The video recording was carried out in a quiet room at 24 °C. The fish were moved from the stocking tank to the observation tank without acclimatisation time and the video recording was started immediately.

### Assay protocol

The acrylic glass observation tank (opaque white, size 36×36×10 cm) is divided into four 18×18 cm quadrants by opaque white acrylic glass dividers. Prior to each assay run (involving the simultaneous assay of four pairs of fish, one pair in each of the four quadrants), the observation tank was rinsed and filled with fresh water (3 cm water depth), and placed inside the test box. Videos were recorded with a Ximea xiQ USB3 colour camera fitted with a 5.5 mm f/1.8 lens mounted above the test tank. The test tank was illuminated with white LED panels providing indirect illumination from above. The front door was closed at the beginning of each run to shield from interference by external stimuli.

Videos were recorded using the Ximea CamTool software (version 4.16), set to capture 30 frames per second. Minor variations from this frame rate, for example 29 frames per second, (movie 20190612_1326_icab_kaga_R) were adjusted for in the downstream analysis. All movies were saved as AVI files. The four reference fishes from the *iCab* line were always introduced to the test tank first and in clockwise order (quadrant order II, I, IV, III), followed by the four test fishes in the same order.

The open field and novel object assay components were carried out consecutively. For the open field assay, as soon as all fish had been introduced in the tank, the fish pairs were recorded for 10 minutes. Subsequently, the door of the test box was opened, and the “novel object” – a small, dark grey, plastic cylinder of 3 cm diameter x 8 cm height was placed in the middle of each quadrant (quadrant order II, I, IV, III). The fish pairs were recorded again for 10 minutes, after which the recording was stopped.

Metadata was recorded for each run including the date, time, tank side, age of reference fish, date on which the reference fish had been used before, test line, test age, frame rate, and additional notes. After each run the door of the tank box was opened, and the fish were removed with a net and placed back into holding containers for transport back to their housing tanks. The test tank was then emptied, and all objects including the quadrant dividers, fish nets, and novel objects were rinsed thoroughly with fresh water, before being returned and refilled for the next run. In most cases, the assay was run concurrently across the two available test tanks (left and right).

To avoid a learning effect, where possible, we only ran the assay on an individual fish once. However, due to limitations in the numbers of *iCab* fish available, of the 77 runs (where a run involved 4 pairs of fish), 27 were performed with *iCab* fishes that had been subjected to the assay once before, and 4 were performed with *iCab* fishes that had been subjected to the assay twice before. When using the same reference fish individuals more than once, we left the maximum amount of time between sessions (at a minimum of 24 hours), and recorded the time and date of any previous sessions. In total, 307 fish pairs were imaged.

### Video pre-processing and tracking

We first pre-processed the videos by splitting them by assay (open field assay in the first 10 minutes and novel object assay in the second 10 minutes) and by quadrant (quadrants I, II, III, and IV), generating 8 separate videos for every 1 original, unprocessed video. Due to slight differences between videos in the appropriate pixel coordinates of the splits, we adjusted them manually for each video. We then tracked the fish in the videos using the open-access software package *idtracker*.*ai* version 5.2.10 (Romero-Ferrero *et al*., 2019). For each video, we manually configured the tracking parameters – such as background subtraction, and minimum and maximum intensity and object areas – to maximise the number of frames that could be successfully tracked. Finally, we labelled in the tracking results each fish as either being the reference fish or test fish, based on which fish had been introduced in the tank first at the start of the video (the reference fish *iCab* was always introduced in the tank first).

For each animal, we calculated the distance and angle travelled between defined time intervals (0.05, 0.08, 0.1, 0.15, 0.2, 0.3, 0.5 and 1 seconds) from their tracking coordinates. In **Figure 1D** we graphically describe how the distance and angle were calculated from a set of 3 consecutive timepoints.

**Figure 1:**
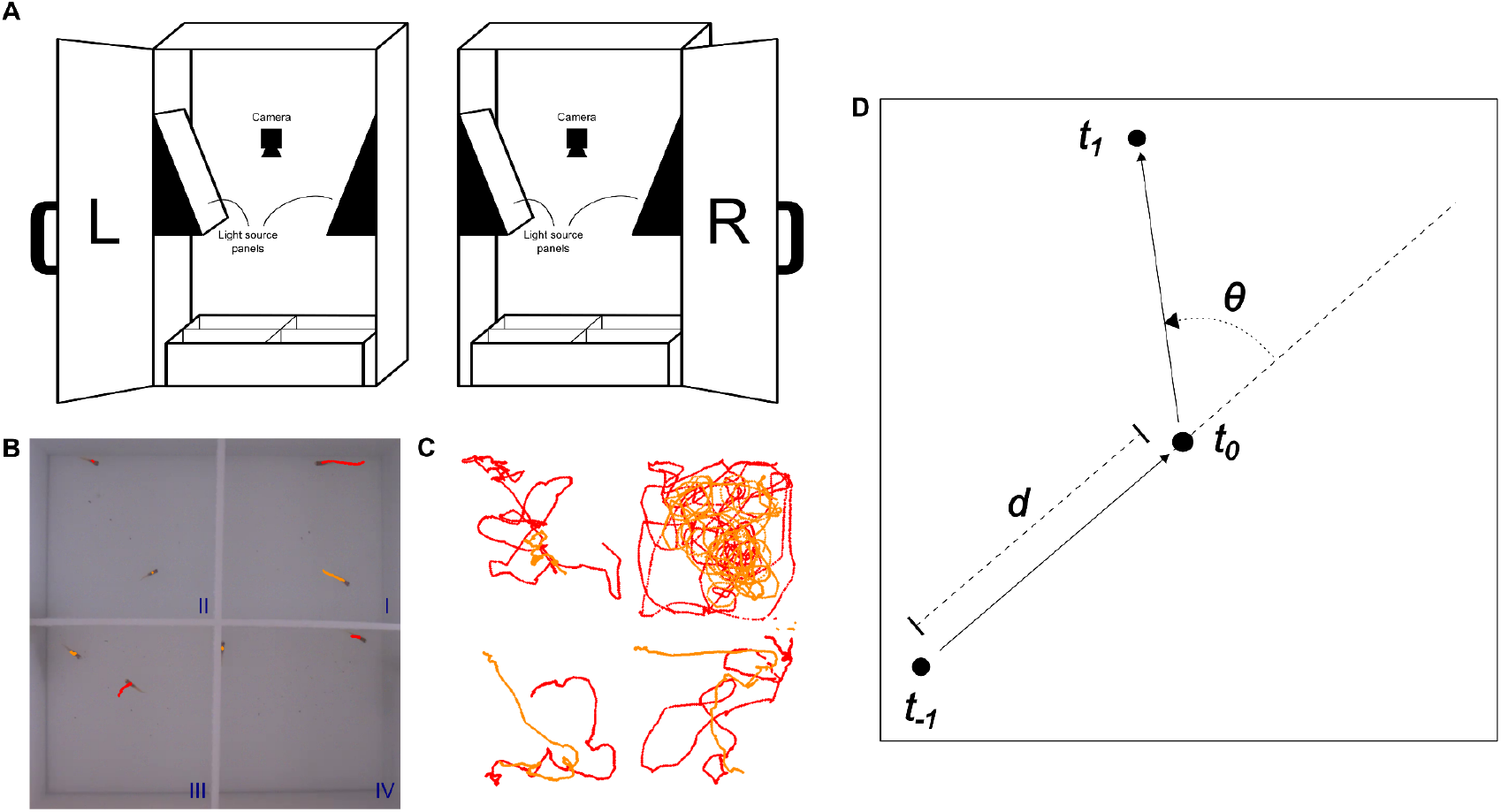
**A**: Experimental setup with two test boxes side-by-side (denoted as “L” for left and “R” for right). Each test box contains one test tank, separated by removable barriers into quadrants, allowing for the simultaneous assaying of four pairs of fish per test tank. The interior of the box is ambiently illuminated by LED lights, and a camera is suspended over the centre of each test tank to record the videos. **B**: Four pairs of fishes in a test tank with labelled quadrants (I, II, III, IV) and strains (red for *iCab*, orange for *HdrR*). **C**: Initial paths of *iCab* reference fish and *HdrR* test fish from the video at panel (B) from 0 to 110 seconds. **D**: The distance of travel (*d*) and the turning angle (θ) were calculated for each fish at each timepoint, and used to train a Hidden Markov Model (see **Methods**). From the fish position at a focal timepoint *t*_0_, the distance was calculated with respect to the previous position *t*_−1_, and the angle was calculated among the *t*_−1_ to *t*_0_ And *t*_0_ to *t*_*1*_ directions of travel.

### HMM parameters

Using the distance and angle measurements as input variables, we trained HMM models with the *hmmlearn* Python package (version 0.3.2, https://github.com/hmmlearn/hmmlearn) across the full dataset with a defined state space (either 5, 10, 12, 14, 15, 16, 17, 18, or 20 states). We sought to determine the combination of parameters that would strike the optimal balance between: a) the ability to detect differences between lines; b) the minimisation of model overfitting; and c) the retention of left-right symmetry between states that would facilitate biological interpretation. To achieve this, we experimented with using different numbers of HMM states (5, 10, 12, 14, 15, 16, 17, 18, 20), and different length of time intervals between which the variables of distance and angle of travel were calculated (0.05, 0.08, 0.1, 0.15, 0.2, 0.3, 0.5 and 1 seconds).

To quantify the level of overfitting by the model, we carried out a two-fold cross-validation process. For each fold, we computed the concordance between the states assigned by an HMM trained on the same fold versus states assigned by an HMM trained on the reciprocal fold (Viterbi paths). We determined that a 0.08-second interval and a state space of 15 yielded the best results, and these settings were used for all downstream analyses. In computing the concordance metric, we matched HMM state identities between the training and validation folds in the way that maximised concordance, starting by matching the most populated training state to the validation state with the highest concordance with it, and proceeding until matching the least populated training state.

### Test for differences in same-state co-occupancy

We tested for strain-specific differences in the frequency of co-occupancy of the same HMM state among test and reference fish pairs. To achieve this, we first calculated the co-occupancy frequency of each possible HMM state pair among the test and reference fish, separately for each assay and fish pair. We then discarded the frequency measurements for non-homologous state pairs (*i*.*e*. we retained only the frequency of co-occupancies in the cases were the test and reference fish occupy the same HMM state), and we ran the Kruskal-Wallis test using the frequency as a dependent variable and the fish strain as a grouping variable. We tested separately for each assay and homologous HMM state pair combination, and corrected our *p*-values for False Discovery Rate.

### Additional software used

All the post-tracking analyses and visualisations were performed using R version 4.3.3 (R Core Team, 2023) the tidyverse suite of R packages (Wickham *et al*., 2019), and cowplot (Wilke, 2020). The code pipeline was constructed with Nextflow version 24.04.2 (Di Tommaso *et al*., 2017).

## Results

### Data collection

Our behavioural assay is 20 minutes long, comprising two consecutively-run 10-minute components: a) an “open field” component, where the fishes are introduced to the test tank and left to swim around freely; and b) a “novel object” component, where a small black plastic cylinder is added to the tank at the beginning of the second 10-minute period, after which the fishes are again left to swim around freely. The assay is run on pairs of fish. Medaka is a seasonal breeder in which photoperiod has a strong effect on physiology and behaviour (López-Olmeda *et al*., 2021), and so in this study we only tested fish that were acclimated to summer conditions. To avoid confounding mating behaviours between males and females, and associated aggressive interactions between males, we used only female fish in all experiments. To increase the throughput of the assay, the test tank was divided into four quadrants with barriers, allowing us to run the assay on four pairs of fish simultaneously. Two test tanks situated side-by-side were used, allowing us to run 8 concurrent assays. The experimental setup that we used is shown in **Figure 1A**.

We assayed a total of 307 pairs of fish, comprising the following counts for each strain pairing: 68 *iCab*/*iCab*, 60 *iCab/HdrR*, 76 *iCab*/*HNI*, 47 *iCab/Kaga*, and 56 *iCab/Ho5*. The fish from the *iCab* strain was denoted as the “reference fish”, and was introduced to the test tank first. The “test fish” was either another *iCab* fish (for the control condition), or a fish from one of the other four strains that were assayed in this experiment (*HdrR, HNI, Kaga*, and *Ho5*). The order in which the strains were assayed across the six days was randomly determined prior to the collection of the data. The test tanks were also rinsed between runs to remove any substances released by subjects during previous runs that could influence the behaviour of the subjects that followed.

### Tracking

Using *idtracker*.*ai* (Romero-Ferrero *et al*., 2019), each individual fish was tracked across at least 78% of frames in each video, with 88% of fish tracked for over 99% of frames. A random selection of 20 videos was reviewed to search for instances of mislabelled fishes due to software errors. We found none, and we therefore concluded that such instances would be absent or very rare.

### Effect of covariates

We examined the effects of several covariates, including date of assay, time of assay, arena quadrant, and test box side (left or right test box). To achieve this, we calculated the mean speed of individuals in *iCab*-*iCab* pairings (*N* = 136) over the course of the entire 20-minute video (including both open field and novel object assay components), and ran an ANOVA test with all the covariates (**Figure 2**).

**Figure 2:**
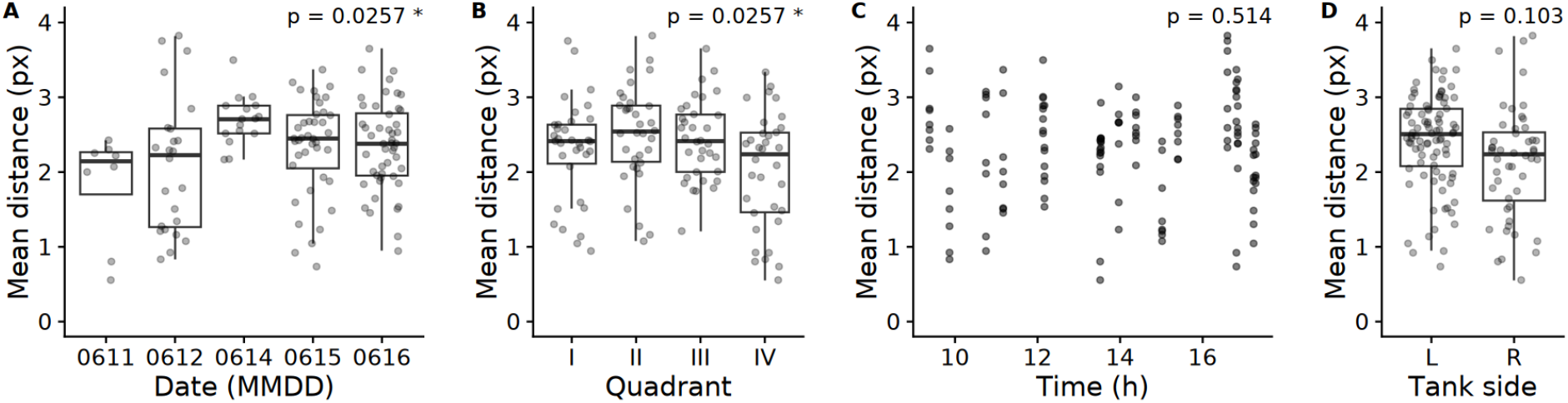
The effect of covariates on the mean distance in pixels travelled by *iCab* individuals in one timestep (0.08 seconds) when paired with another *iCab* over the course of the full video (including both open field and novel object assays). *p-*values were calculated from an ANOVA test with all four covariates included as terms, and were adjusted for false discovery rate.

We found some significant differences for the covariates date of assay and tank quadrant (*p* = 0.0254 and *p* = 0.0254), but not for time of assay or test box side. Given this significant result, we decided to use all the covariates in the downstream analysis to compensate for the experimental variation. We show the regions of the test tank that were most frequently occupied by the fish stratified by quadrant, tank, and assay in **Figure S1**.

### Choice of time interval and number of HMM states

To determine the optimal parameters for the HMM’s classification of behaviours, we sought to reduce overfitting (which tends to favour using a lower number of HMM states) while maximising the ability to distinguish between strains based on the relative time they spent in each HMM state (which tends to favour a higher number of HMM states). We additionally considered the time interval within which the distance and angle of travel were measured. We also wanted to avoid HMMs with large asymmetries (for example an HMM with a left-turning state but no right-turning state) because of the difficulties that they would pose in interpreting the biological meaning of those asymmetric states. **Figure 3** sets out the comparison of HMM parameters on two measures designed to quantify, respectively, the level of overfitting (mean concordance), and the quantification of differences between strains (summed Kruskal-Wallis χ^2^) (see **Methods**).

**Figure 3:**
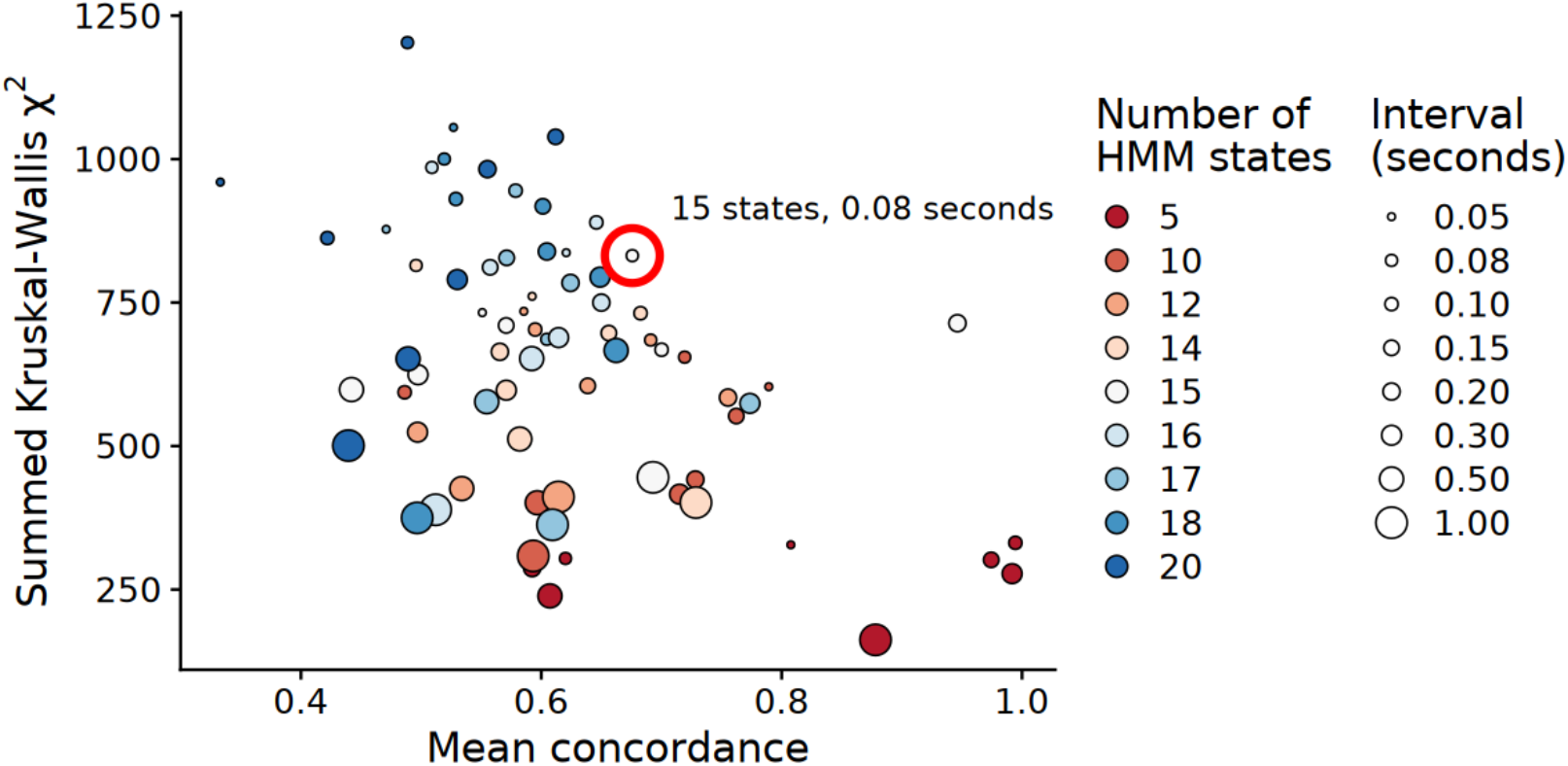
Selection of the number of hidden states used in the HMM and of the time interval in which distance and angle of travel are calculated. Horizontal axis: Mean concordance between states assigned by HMMs in a 2-fold cross-validation procedure. Vertical axis: Kruskal-Wallis test statistic comparing strains based on the proportion of time spent in each HMM state, summed across all states. The size of the points represents the interval, in seconds, between which the distance and angle of travel were calculated. The colour indicates the number of hidden states used in the HMM. The circled parameter combination was used for further analysis.

Based on these results and on the visualisation of the polar plots for each combination of state number and time interval, we selected the combination of 15 hidden states with a 0.08-seconds time interval, because out of the remaining combinations it appeared to optimally balance the level of overfitting, the detection of differences between strains, and the absence of asymmetric states. We also considered the combination of 15 hidden states and 0.2-seconds time interval, which had high concordance and only moderately lower summed Kruskal-Wallis statistic. However, the state space in this combination was less descriptive, with no left or right turning states, and for this reason we preferred the 15-states and 0.08-seconds time interval combination. The distances and angles of travel for the selected HMM are shown in **Figure 4A**.

**Figure 4:**
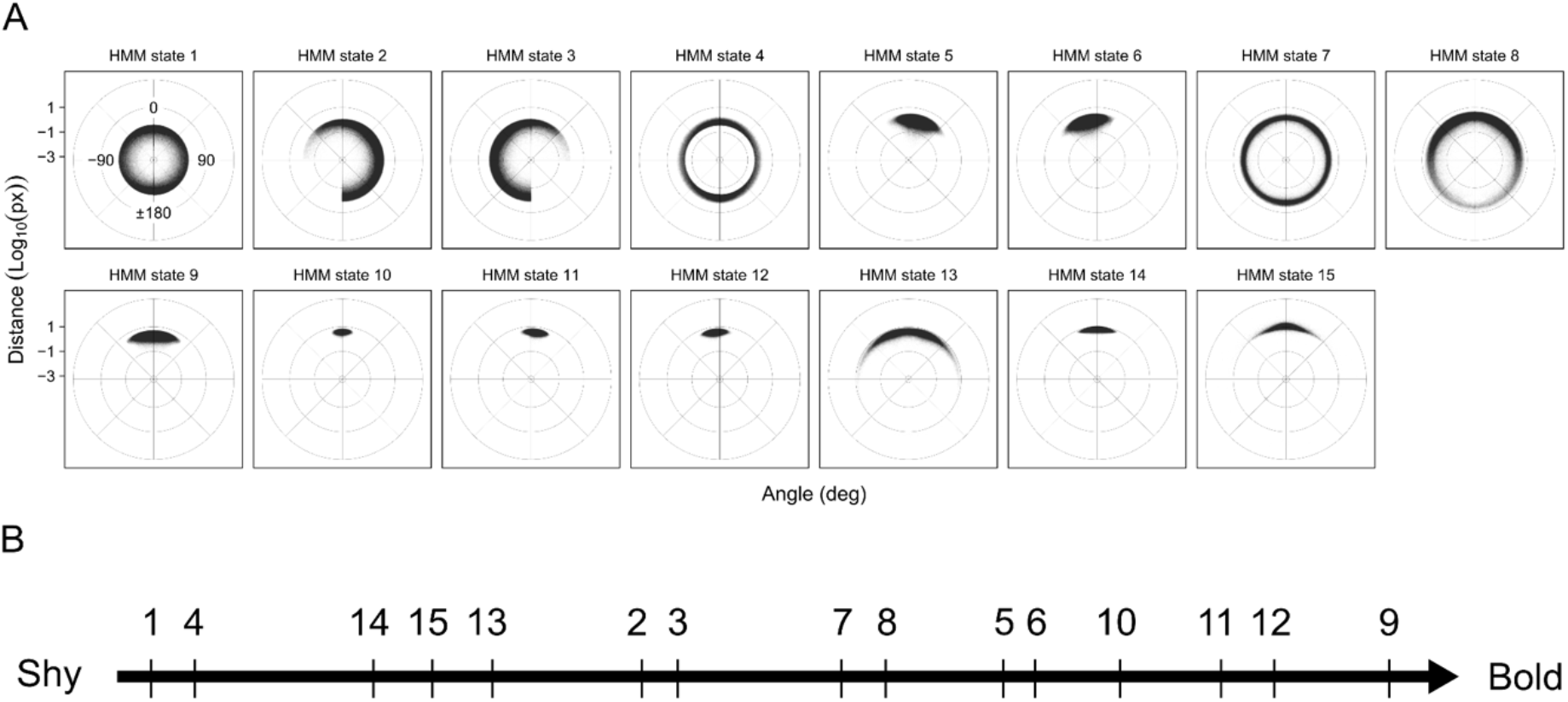
**A**: Classification of movements by the 15-state HMM, based on distance (in log_10_(pixels)) and angle of travel in a time interval of 0.08 seconds. States are sorted in ascending order by mean distance of travel. Each point represents the radial distance in log10(pixels) that a fish travelled from its previous to current location, and the polar angle between the incoming and outgoing directions of travel (see **Figure 1D** for a graphical representation). To illustrate, a point at 45° far from the pole represents a fast, forward movement to the right, and a point at -135° close to the pole represents a slow, backward movement to the left. **B**: Tentative positioning of the HMM states along the bold-shy behavioural axis (see main text and **Supplementary note**).

### Positioning of the HMM states along the bold-shy behavioural axis

We tentatively ordered the HMM states along the bold-shy behavioural axis (**Figure 4B**) by considering the distribution of locomotion angle and distance, position within the tank and frequency of occupancy during the open field test (**Figure S1 and S2**). Temporal frequency during the open field test was given the highest weight, as a decrease in occupancy over time can be assumed to rely on habituation (Lucon-Xiccato, Loosli, *et al*., 2022). We then considered tank position, where a high density of occupancy along the wall hints at thigmotaxis-related avoidance of more central tank regions, indicating shyness (Lucon-Xiccato, Loosli, *et al*., 2022). Finally, initial slow locomotion was considered to indicate fear-related freezing behaviour, whereas fast forward swimming at later time points was considered to indicate bold exploratory behaviour. We provide state-specific details of the rationale for our HMM state ordering in the **Supplementary Notes**.

### Direct genetic effects

To determine whether the test fish strains differed in the proportions of time they spent in each state, we ran a separate ANOVA test for each combination of assay component (open field or novel object) and state (states 1 to 15). The proportion of time each individual fish spent within a state was first inverse-normalised within each combination of assay and state. The date of assay, time of assay, tank quadrant, and tank side were included as covariates. The inverse normalisation ensures that the frequency metric has a gaussian distribution, and so the ANOVA test is valid. For the inverse normalisation we followed the procedure described in Yang *et al*., 2012. *p*-values were adjusted for the False Discovery Rate (FDR) using the Benjamini & Hochberg correction (Benjamini and Hochberg, 1995) implemented in the p.adjust function in R. We use letter *q* to represent the FDR adjusted metric.

The results of this analysis (*p*-values and variance explained) are set out in **Table S1**. The test fish strains differed significantly in the proportion of time spent in a given state (*q* < 0.05, FDR-adjusted) for 11 out of 15 states in the open field assay (1.17 ∗ 10^−24^ ⩽ q ⩽ 0.0236), and 8 out of 15 states for the novel object assay (2.79 ∗ 10^−13^ ⩽ q ⩽ 0.000156), with the strain of the test fish explaining up to 29.7% of the variance in the proportion of time spent in a given state. For some states, as expected from our initial analysis on speed, there was also a significant difference between quadrants and dates of assay (open field: 2.81 ∗ 10^−7^ ⩽ q ⩽ 0.899; novel object: 4.42 ∗ 10^−8^ ⩽ q ⩽ 0.968).

In **Figure 5** we depict the time dependence of HMM states over the course of the video, and the regions of the test tank that were most frequently occupied by the different strains. All the strains show a predominance of slow-moving states (i.e. states 1 to 4, dark purple, to light blue) at the beginning of each assay component. Northern medaka strains (*Kaga, HNI*) differ from southern medaka strains (*iCab, HdrR*, and *Ho5*) for presenting a smaller proportion of slow-moving states at the beginning of the open field assay. On the contrary, the later stages of the open field assay appear to be more uniform in state usage across strains. Differently from other strains, *Kaga* tends to spend more time in the fast and forward-moving state 15 (yellow) at the very beginning of the open field assay than at other times.

**Figure 5:**
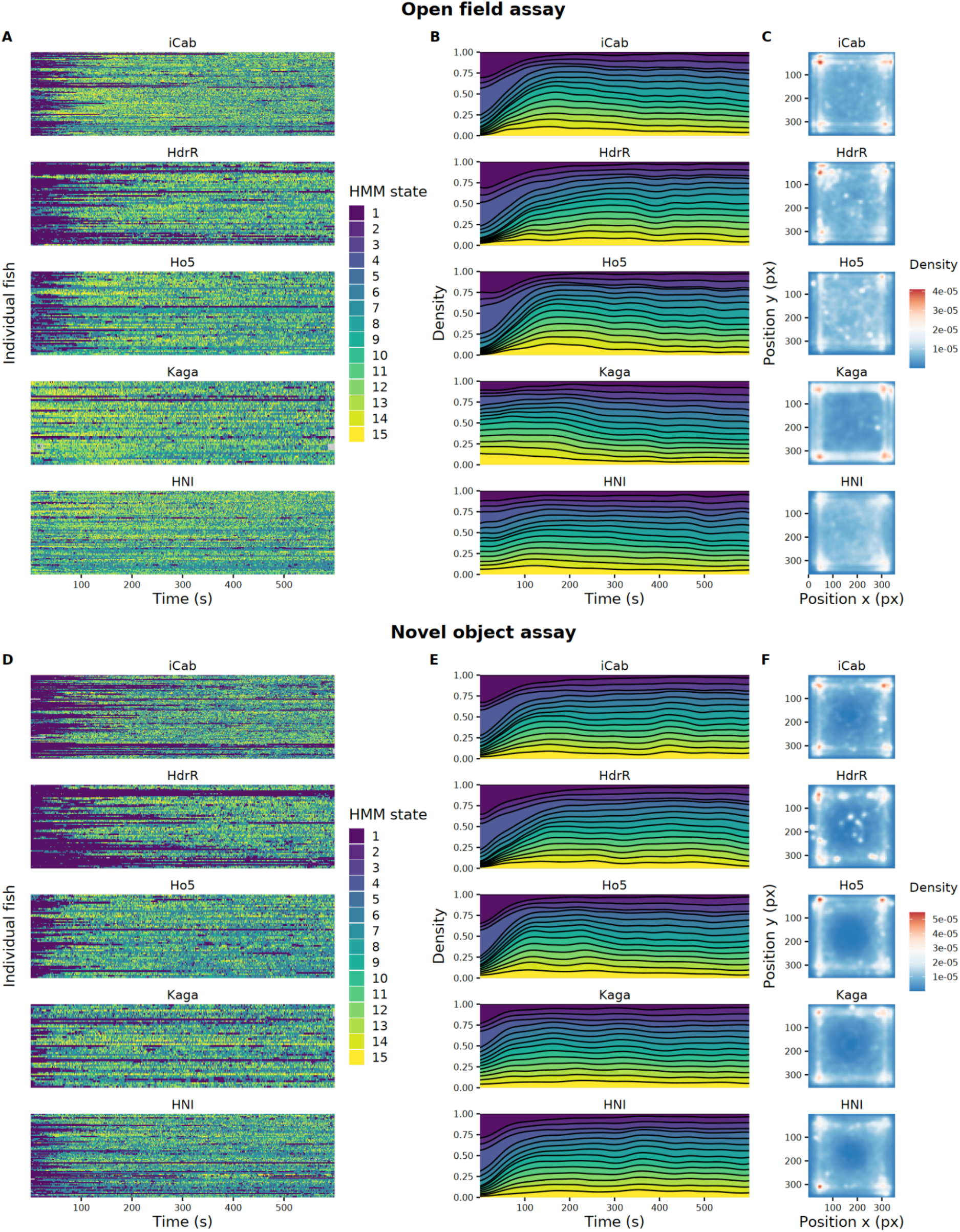
Differences between test fish strains in the HMM states they occupy and their position in the arenas during the open field (panels A, B, C) and novel object (panels D, E, F) assay components. The HMM states are ordered by average distance travelled (same encoding as in **Figure 4**). **A, D**: Transitions between HMM states across time for each individual test fish, grouped by strain. **B, E**: Time-dependent density of usage of HMM states across samples, separated by strain. **C, F**: Densities of the test tank locations occupied by each strain throughout the assay. The y axis is inverted because pixel positions are measured from top to bottom.

### Social genetic effects

To determine whether the *iCab* reference fish altered their behaviour depending on the inbred strain of their tank partner, we applied the same analysis and model as above using only from the *iCab* reference fish instead than from the test fish itself. The results for this analysis are set out in **Table S2**.

Similarly to what we observed for direct genetic effects, in the novel object assay we notice a difference among northern and southern strains, with northern strains showing lower occupancy of slow-moving states at the beginning of the videos. The *iCab* reference fish differed significantly in the proportion of time they spent in a given state depending on the strain of their tank partner (*q* < 0.05, FDR-adjusted) for 2 out of 15 states in the open field assay (1.58 ∗ 10^−5^ ⩽ q ⩽ 2.99 ∗ 10^−5^), and 3 out of 15 states in the novel object assay (1.90 ∗ 10^−5^ ⩽ q ⩽ 1.54 ∗ 10^−2^). The strain of the tank partner explained up to 8.64% of the variance in the proportion of time the *iCab* reference spent in a given state.

We observe the behavioural patterns of the *iCab* reference fish tend to reflect those of the test fish strains they are paired with (**Figure 6**). The *iCab* reference fish spend less time in the slower-moving states 1 to 4 when in the presence of the faster-moving northern Japanese strains *HNI* and *Kaga* (19.8% to 22.6% of the time). On the contrary, the reference fish spends more time in slower-moving states when paired with the southern strains *iCab* and *HdrR* (25.0% and 28.0% of the time). Similarly to what the *Kaga* test fish themselves do, at the start of the open field assay the reference fish paired with *Kaga* show an increased occupation of the fast-moving state 15.

**Figure 6:**
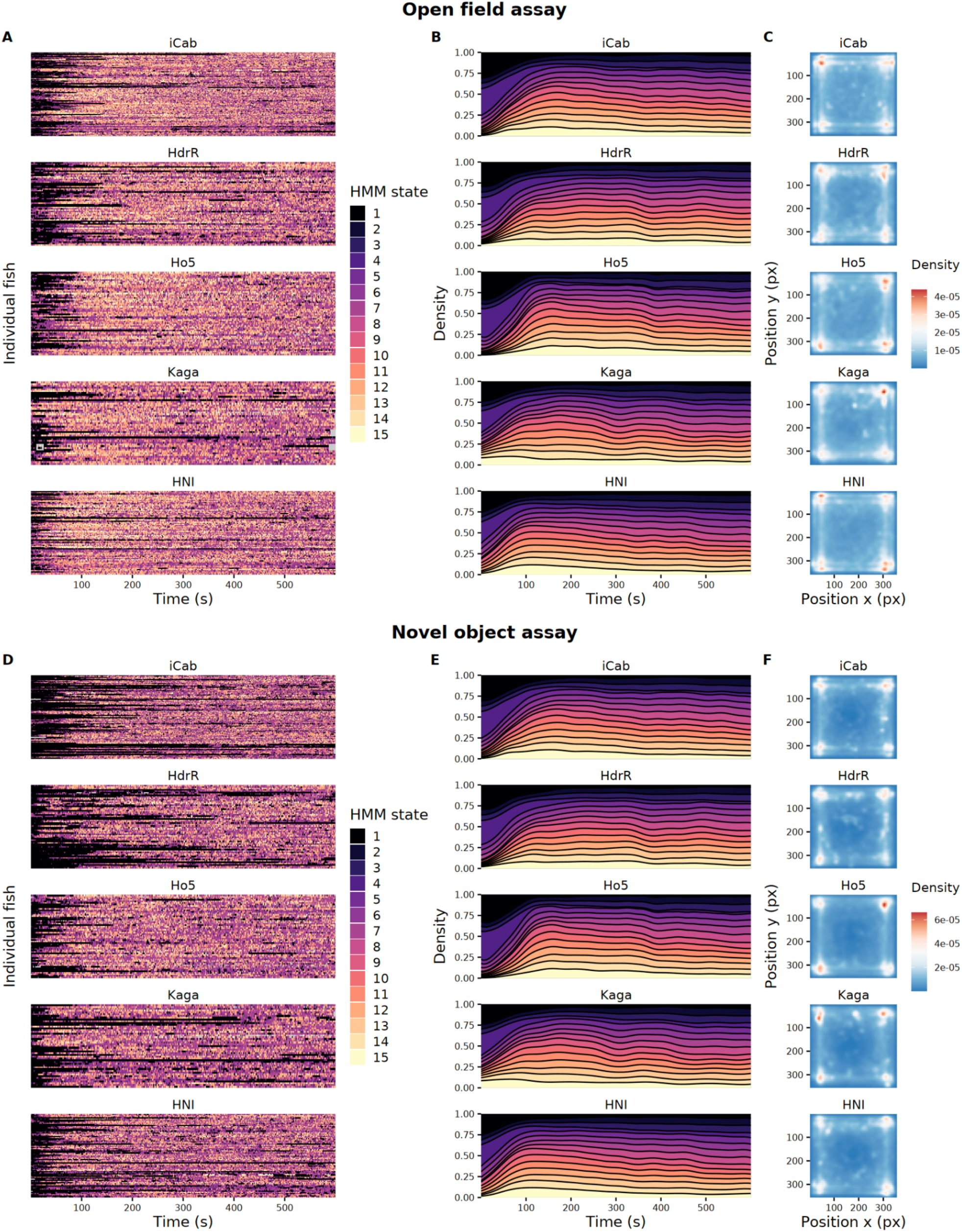
Differences in the HMM states and location in the test arenas occupied by the iCab reference fish depending on the test fish strain it was paired with. The HMM states are ordered by average distance travelled (same encoding as in **Figure 4**). The results are presented separately for the open field (panels A, B, C) and novel object (panels D, E, F) assay components. **A, D**: Transitions between HMM states across time for each individual reference fish, grouped by strain of the tank partner. **B, E**: Time-dependent density of usage of HMM states across reference samples, separated by strain of the tank partner. **C, F**: Densities of the test tank locations occupied by the reference fish throughout the assays depending on the strain of the tank partner. The y axis is inverted because pixel positions are measured from top to bottom.

To better quantify the degree to which the *iCab* reference fish behaviour is influenced by the strain of its tank partner, we calculated the co-occupancy among HMM states occupied by the test and reference fish, stratified by assay component and test fish strain (**Figure 7**). For each combination of assay component and state, we ran a Kruskal-Wallis test (see **Methods**) to determine whether there were differences in the frequencies of same-state co-occupancy under different strain pairings (*q* < 0.05, FDR-adjusted). We observe significant differences in 7 out of 15 HMM states in the open field assay, and 5 out of 15 HMM states in the novel object assay. We report these results in **Table S3**.

**Figure 7:**
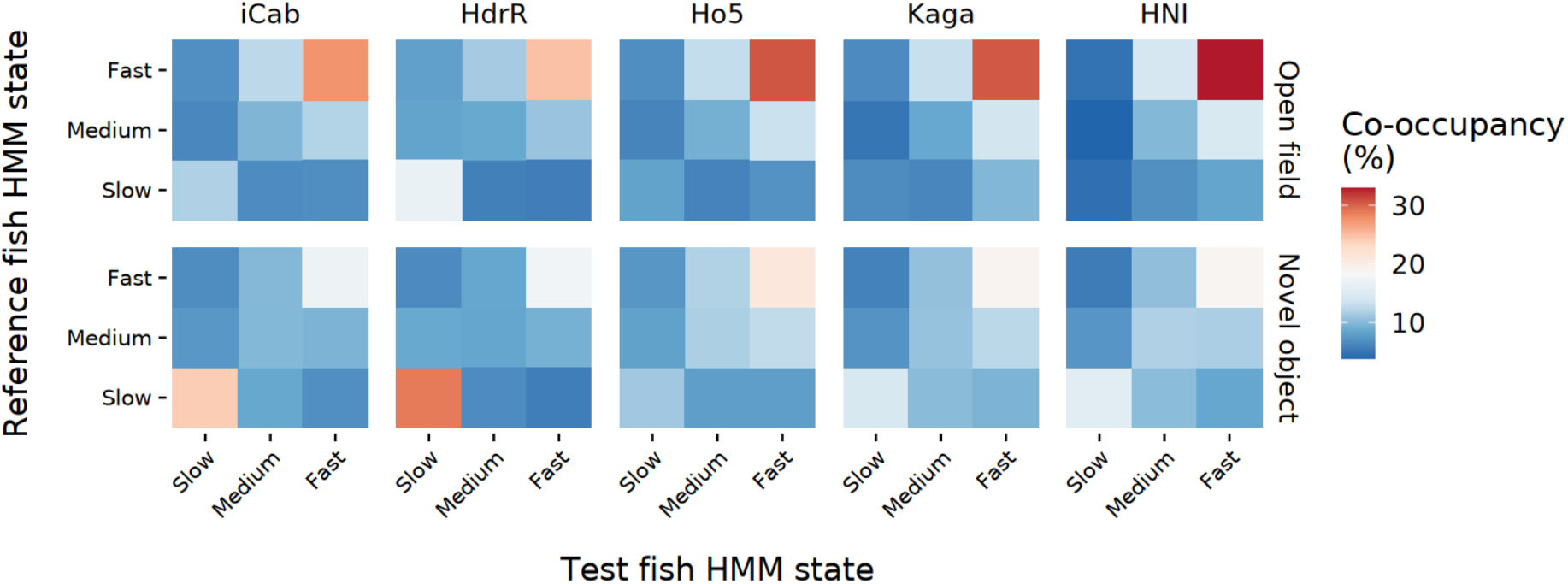
Frequency of HMM state co-occupancy between test and reference fish, calculated across all videos per strain-pairing and assay component. For ease of visualisation, slow (1-4), medium (5-8), and fast (9-15) HMM states are grouped together.

## Discussion

In this study we have described a robust assay for measuring bold-shy behaviours in medaka fish that can reliably detect differences: (a) between individuals from different inbred strains, allowing for the quantification of direct genetic effects on behaviour; and (b) between the behaviour of individuals when paired with tank partners from a different strain allowing for the quantification of social genetic effects. We show that Hidden Markov Models (HMMs) allow for a reasonable classification of medaka behavioural modes based on the direction and angle of travel within set time intervals. In principle, this method can be expanded to include additional behavioural features such as proximity to the wall or other objects, inwards/outwards orientation, proximity to the tank partner, and other metrics related to leader-follower dynamics (a possibility that will be tested in future studies). Alternative methods to HMMs could be used, for example, recurrent neural networks (Yu *et al*., 2019) or transformer-based models (Vaswani *et al*.). However, a benefit of the simpler HMM framework is that the states are more interpretable and can easily be integrated into established statistical and genetic frameworks, as we have done in this work.

We note that the slower moving states exhibit higher relative levels of noise because even when a fish is almost completely still, the fish object’s centroid (upon which the variables of distance and angle of travel are calculated) will tend to move by one or several pixels through minor changes in the segmentation of the object. We also found that behaviours exhibited during the assay show some variance across the covariates date of assay, time of assay, tank quadrant, and tank side.

We reported that inbred strains of medaka fish can be distinguished by the proportion of time they spend in certain states. By design, this experiment had enough replicates of each genetic pairing in each potential quadrant or tank so that many sources of variation could be detected; this led to highly significant FDR-corrected genetic effects, in some cases with substantial proportion of variance explained. As well as the overall ability to show genetic effects, we were also able to dissect some features of medaka bold-shy behaviour. For example, the slowest states (1 to 4), capturing no or minimal movement, were the states that most clearly separated the strains, and these differences were most evident at the beginning of the open field and novel object assays. The southern medaka strains (*iCab, HdrR*, and *Ho5*) spent significantly more time in the slower moving states at the beginning of each assay component relative to the northern medaka strains (*Kaga* and *HNI*). This is consistent with the hypothesis that for southern Japanese strains, the “freeze” reflex is caused by anxiety, which eventually dissipates over time during habituation. Similar observations of initial anxiety represented by freezing behaviour and subsequent habituation have been made in other studies using open field assays with *iCab* fish (Lucon-Xiccato, Loosli, *et al*., 2022). On the other hand, the northern Japanese strains *Kaga* and *HNI* spent little time in the slow-moving states at the beginning of the video, which indicates either that their habituation sets in earlier, or that their stress and anxiety is expressed to a lesser degree as freezing behaviour. The latter appears to be more likely for *Kaga*, as in the open field assay, it spends more time in the faster-moving states at the beginning of the video (especially state 15) and then slows down thereafter, which suggests that its higher level of movement may be induced by stress. It is interesting then that once the novel object is introduced, *Kaga* tends to move slowly like the other strains. Obviously, its introduction to a novel environment and exposure to a potentially dangerous object elicits different behavioural responses in this strain. We observe strong thigmotaxis behaviour (*e*.*g*. moving along the sides of the tank) across all fish strains, and this behaviour is particularly pronounced for the Kaga strain in the open field assay.

With respect to social genetic effects, we observed that a fish’s behaviour is affected by the differential behaviour of its tank partner, although the effect is less powerful than the direct genetic effect on a fish’s own behaviour. These social genetic effects are detectable when observing the proportions of time the reference fishes spend in certain states over the course of the video, and can also be investigated by observing the frequency of state co-occupancy among tank partners. Interestingly, there are more significant changes in state frequency for social genetic effects in the novel object setting. The sudden introduction of a novel object in the tank may be a stronger stressor than being placed in an unfamiliar environment (open field assay), and as such a social animal such as medaka may in this setting rely more heavily on the presence of a partner for cues. In this study we only used one strain as the reference fish, but future experiments can expand on this analysis by using different strains as the reference, thereby exploring how a strain’s genetics influence the degree of their behavioural plasticity, and how the distinct behaviours of strains may interact.

In summary we show that our experimental setup in combination with the *idtracker*.*ai* movement tracking and the HMM analysis can reliably detect both direct and social genetic effects. This creates the opportunity to carry out a similar study on a larger panel of inbred strains, such as the MIKK panel (Fitzgerald *et al*., 2022). Expanding our current framework of exploring strain-specific differences, we plan to perform F2 crosses among different inbred strains with the aim of identifying genetic variants associated with differences both in the fish’s own behaviour, and also in the degree to which an individual transmits their behaviour onto their social companions. This, in turn, will shed light on longstanding biological questions concerning direct and indirect influences on behaviour, their physiological bases, and their adaptive purposes.

## Acknowledgements

The authors acknowledge Nadeshda Wolf and Natalja Kusminski for excellent fish husbandry and Daniel Marcato for technical support. This research was supported by the European Research Council Synergy Grant IndiGene [grant number 810172].

## Data and code availability

All code used for the analysis is available on GitHub: https://github.com/birneylab/medaka_behaviour_pilot. The data underlying this article are available on the EBI Bioimage Archive: https://doi.org/10.6019/S-BIAD1421.

