## Supplementary material for "Measurement and classification of bold-shy behaviours in medaka fish"

### Supplementary Notes

#### Positioning of the HMM states along the bold-shy behavioural axis

In the main test we describe the general principles that we followed in positioning HMM states along the bold-shy behavioural axis. Using these criteria, we conclude that HMM states 1 and 4 indicate shy behaviour as they are more frequent at the beginning of the videos (both in the open field as well as the novel object tests) and are associated with low levels of locomotion (**Figure S1** and **S2**). State 9 indicates a bold state as its frequency increases over time and is associated with high levels of locomotion encompassing the entire tank area. Similarly states 5, 6 and 10-12 show high locomotion activity in all tank areas, though with a less pronounced temporal increase of frequency. States 13-15 are more ambiguous in that they are associated with high locomotion and a slight temporal increase. However, tank positions are mostly at the perimeter. Furthermore, the frequency reaches an early peak at about 100 seconds followed by a gradual decrease. This is reminiscent of the behavioural pattern observed with Kaga in the open field test where initially high locomotion activity subsequently decreases (**Figure 5**). We therefore conclude that this state can indicate stress related high activity. The occupancy of state 7 and 8 increases over time and is associated with moderate locomotion and high occupancy of corner regions, suggesting exploratory behaviour of these regions. Finally states 2 and 3 exhibit slow locomotion and corner occupancy which could indicate a cautious resting state.

### Supplementary Tables

| Assay component | HMM State | p-value (FDR-adjusted) | Significance | Variance explained |
|---|---|---|---|---|
| Open field | 1 | 1.17e-24 | **** | 0.297 |
| Open field | 2 | 4.75e-07 | **** | 0.101 |
| Open field | 3 | 6.42e-07 | **** | 0.0977 |
| Open field | 4 | 6.06e-21 | **** | 0.261 |
| Open field | 5 | 2.36e-02 | * | 0.0358 |
| Open field | 6 | 1.73e-02 | * | 0.0384 |
| Open field | 7 | 5.54e-07 | **** | 0.0964 |
| Open field | 8 | 3.76e-05 | **** | 0.0699 |
| Open field | 9 | 8.54e-05 | **** | 0.0673 |
| Open field | 10 | 3.80e-01 | ns | 0.0161 |
| Open field | 11 | 2.36e-01 | ns | 0.0209 |
| Open field | 12 | 3.11e-01 | ns | 0.0185 |
| Open field | 13 | 5.35e-03 | ** | 0.0452 |
| Open field | 14 | 2.01e-02 | * | 0.0406 |
| Open field | 15 | 1.98e-01 | ns | 0.0221 |
| Novel object | 1 | 2.79e-13 | **** | 0.175 |
| Novel object | 2 | 1.53e-07 | **** | 0.109 |
| Novel object | 3 | 2.25e-07 | **** | 0.106 |
| Novel object | 4 | 5.85e-10 | **** | 0.139 |
| Novel object | 5 | 1.48e-04 | *** | 0.0681 |
| Novel object | 6 | 1.56e-04 | *** | 0.0681 |
| Novel object | 7 | 2.25e-07 | **** | 0.103 |
| Novel object | 8 | 7.52e-01 | ns | 0.00636 |
| Novel object | 9 | 2.01e-06 | **** | 0.0944 |
| Novel object | 10 | 2.93e-01 | ns | 0.019 |
| Novel object | 11 | 3.87e-01 | ns | 0.0154 |
| Novel object | 12 | 4.75e-01 | ns | 0.0135 |
| Novel object | 13 | 3.62e-01 | ns | 0.0148 |
| Novel object | 14 | 8.68e-01 | ns | 0.00513 |
| Novel object | 15 | 6.97e-01 | ns | 0.00938 |

**Table S1**: Strain-dependent differences and variance explained from the ANOVA test for the proportion of time spent in each HMM state by the test fish (direct genetic effect) across test fish strains for the open field and novel object assay components.

| Assay component | HMM State | p-value (FDR-adjusted) | Significance | Variance explained |
|---|---|---|---|---|
| Open field | 1 | 2.99e-05 | **** | 0.0815 |
| Open field | 2 | 8.81e-01 | ns | 0.00361 |
| Open field | 3 | 8.91e-01 | ns | 0.0031 |
| Open field | 4 | 1.58e-05 | **** | 0.0864 |
| Open field | 5 | 4.74e-01 | ns | 0.0102 |
| Open field | 6 | 4.45e-01 | ns | 0.0108 |
| Open field | 7 | 6.82e-01 | ns | 0.00696 |
| Open field | 8 | 2.40e-01 | ns | 0.0169 |
| Open field | 9 | 6.74e-01 | ns | 0.00709 |
| Open field | 10 | 2.02e-01 | ns | 0.021 |
| Open field | 11 | 1.14e-01 | ns | 0.0269 |
| Open field | 12 | 9.43e-02 | ns | 0.0285 |
| Open field | 13 | 1.29e-01 | ns | 0.0233 |
| Open field | 14 | 1.60e-01 | ns | 0.0244 |
| Open field | 15 | 6.85e-01 | ns | 0.00719 |
| Novel object | 1 | 3.07e-05 | **** | 0.0774 |
| Novel object | 2 | 1.15e-01 | ns | 0.0265 |
| Novel object | 3 | 1.15e-01 | ns | 0.0263 |
| Novel object | 4 | 1.90e-05 | **** | 0.0815 |
| Novel object | 5 | 2.88e-01 | ns | 0.0165 |
| Novel object | 6 | 2.53e-01 | ns | 0.018 |
| Novel object | 7 | 1.54e-02 | * | 0.0393 |
| Novel object | 8 | 5.62e-02 | ns | 0.0292 |
| Novel object | 9 | 2.35e-01 | ns | 0.0203 |
| Novel object | 10 | 2.40e-01 | ns | 0.0199 |
| Novel object | 11 | 1.60e-01 | ns | 0.0246 |
| Novel object | 12 | 1.62e-01 | ns | 0.0243 |
| Novel object | 13 | 2.31e-01 | ns | 0.0182 |
| Novel object | 14 | 2.38e-01 | ns | 0.0191 |
| Novel object | 15 | 4.26e-01 | ns | 0.0138 |

**Table S2**: Strain-dependent differences and variance explained from the ANOVA test for the proportion of time spent in each HMM state by the reference fish (social genetic effect) across test fish strains for the open field and novel object assay components.

| Assay component | HMM State | p-value (FDR-adjusted) | Significance |
|---|---|---|---|
| Open field | 1 | 1.68e-07 | **** |
| Open field | 4 | 7.98e-06 | **** |
| Open field | 7 | 1.30e-02 | * |
| Open field | 2 | 1.30e-02 | * |
| Open field | 3 | 8.03e-03 | ** |
| Open field | 6 | 5.42e-01 | ns |
| Open field | 5 | 6.85e-01 | ns |
| Open field | 8 | 2.82e-01 | ns |
| Open field | 9 | 3.69e-01 | ns |
| Open field | 12 | 3.07e-01 | ns |
| Open field | 11 | 2.82e-01 | ns |
| Open field | 10 | 4.83e-02 | * |
| Open field | 13 | 7.30e-01 | ns |
| Open field | 14 | 7.83e-01 | ns |
| Open field | 15 | 1.30e-02 | * |
| Novel object | 1 | 2.54e-07 | **** |
| Novel object | 4 | 4.79e-07 | **** |
| Novel object | 7 | 8.03e-03 | ** |
| Novel object | 2 | 1.30e-02 | * |
| Novel object | 3 | 4.83e-02 | * |
| Novel object | 6 | 9.10e-02 | ns |
| Novel object | 5 | 9.10e-02 | ns |
| Novel object | 8 | 7.76e-01 | ns |
| Novel object | 9 | 9.10e-02 | ns |
| Novel object | 12 | 7.54e-01 | ns |
| Novel object | 11 | 7.54e-01 | ns |
| Novel object | 10 | 8.80e-01 | ns |
| Novel object | 13 | 7.76e-01 | ns |
| Novel object | 14 | 5.66e-01 | ns |
| Novel object | 15 | 3.64e-01 | ns |

**Table S3**: Strain-dependent differences (Kruskal-Wallis test) in the co-occupancy of the same HMM state by the test and reference fish, stratified by assay component and HMM state.

### Supplementary Figures

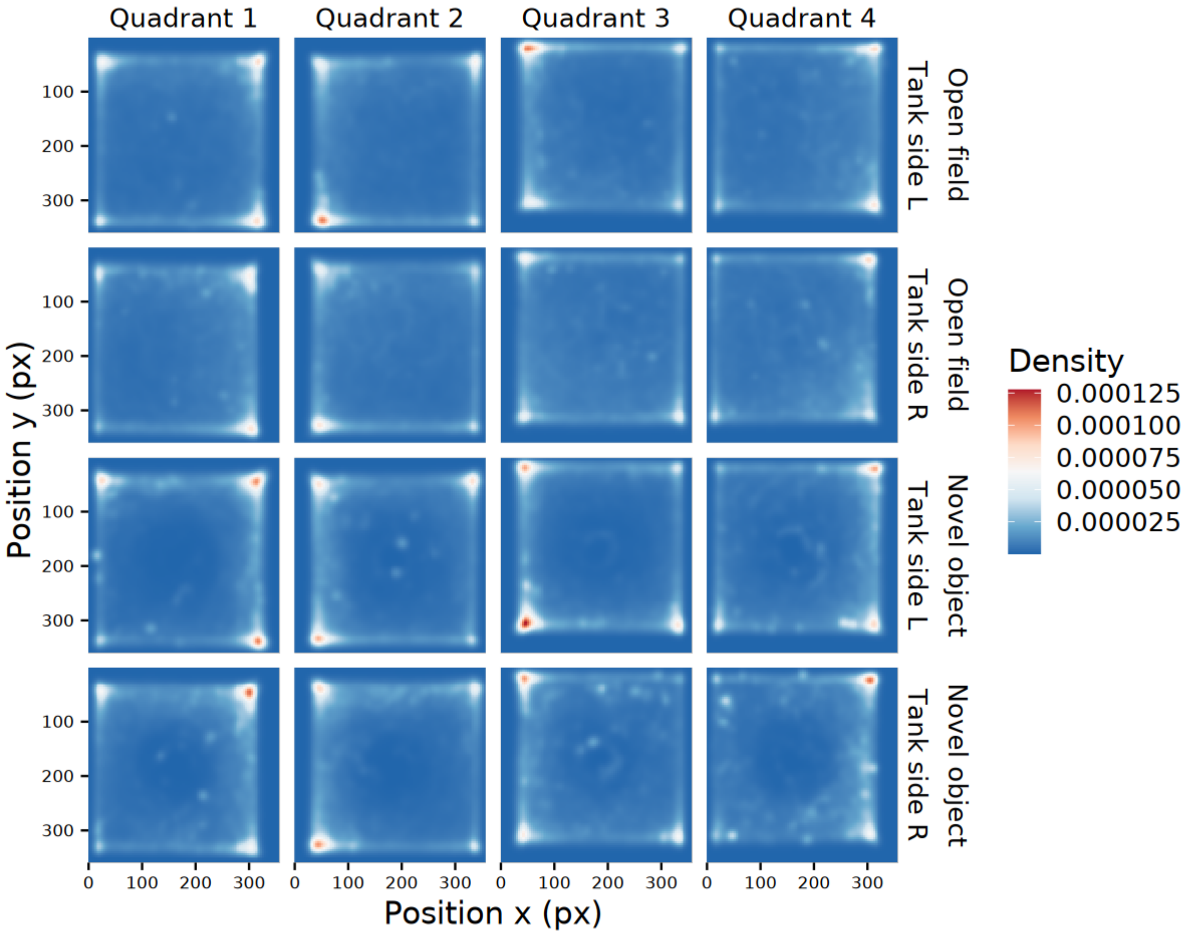


**Figure S1:** Areas of the test tanks occupied by the fishes stratified by assay component, quadrant, and tank side (left or right testing apparatus).

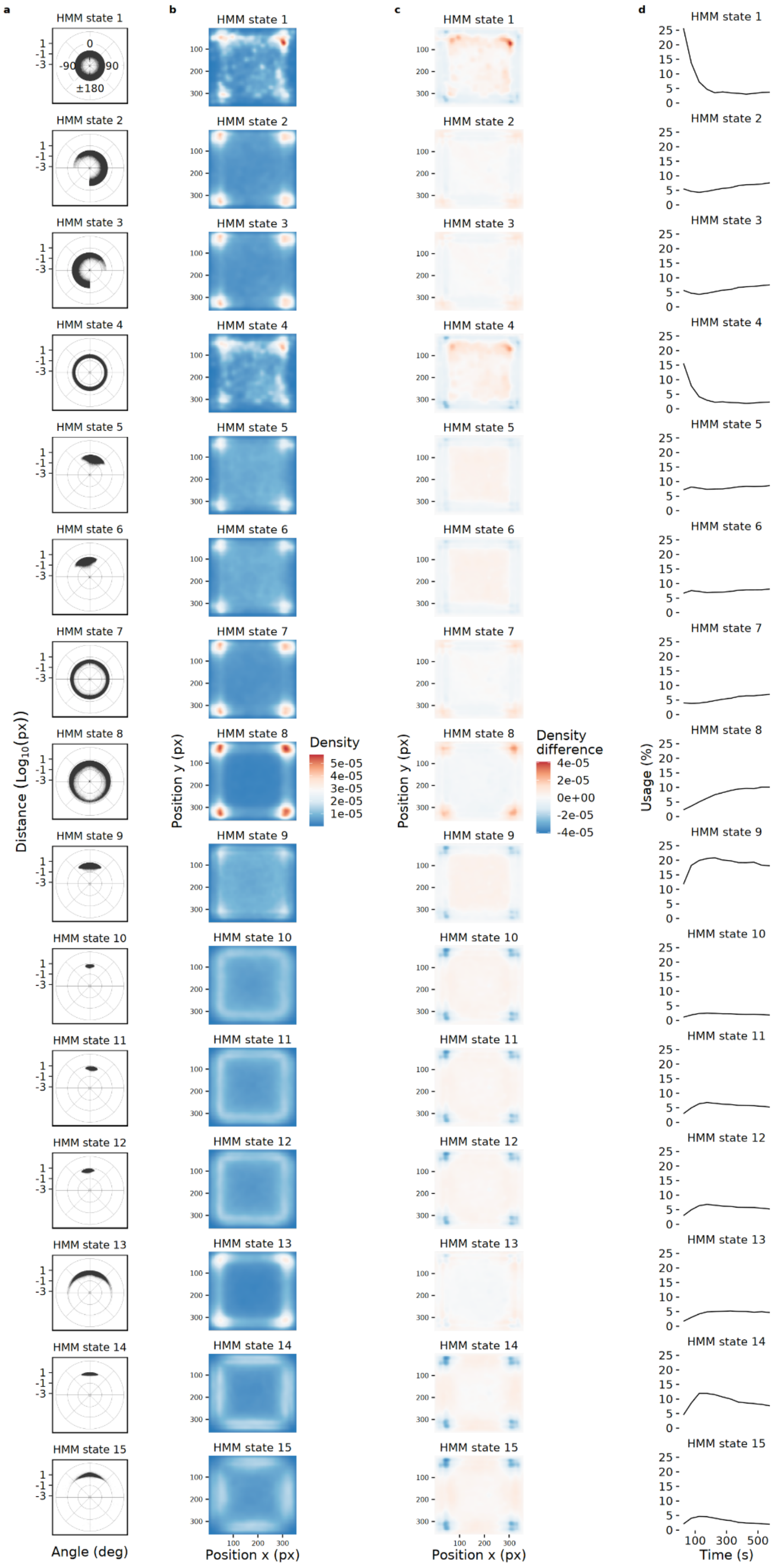


**Figure S2:** Characterization of the hidden states in the open field assay component with respect to distances and angles (a), position of the fish in the tank (b), difference in the density of fish positions in the tank with respect to the fish position density calculated excluding the first 400 seconds of habituation (c), and hidden state usage across time (d).

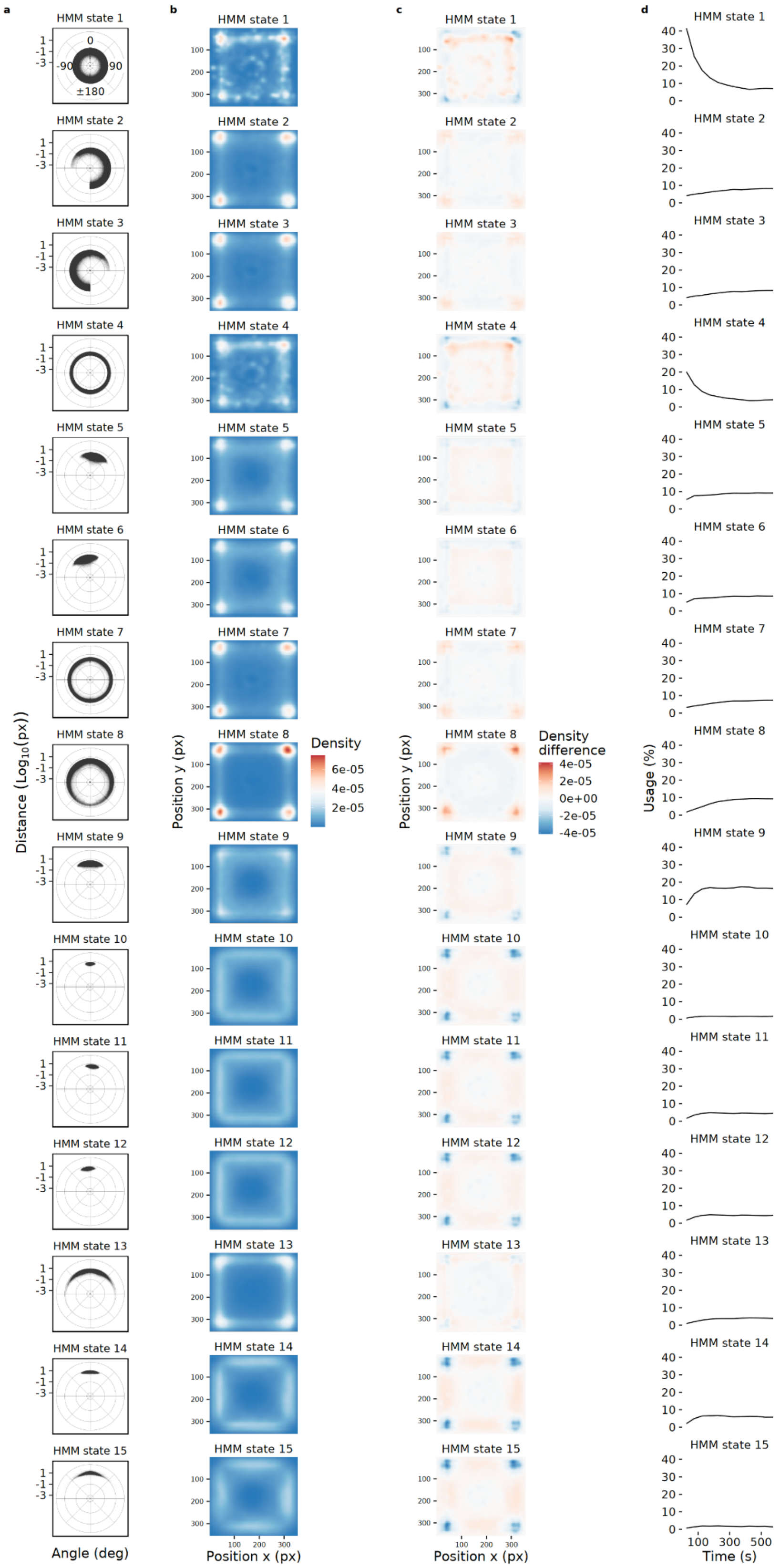


**Figure S3:** Characterization of the hidden states in the novel object assay component with respect to distances and angles (a), position of the fish in the tank (b), difference in the density of fish positions in the tank with respect to the fish position density calculated excluding the first 400 seconds of habituation (c), and hidden state usage across time (d).
